# Kallikrein 13: a new player in coronaviral infections

**DOI:** 10.1101/2020.03.01.971499

**Authors:** Aleksandra Milewska, Katherine Falkowski, Magdalena Kalinska, Ewa Bielecka, Antonina Naskalska, Pawel Mak, Adam Lesner, Marek Ochman, Maciej Urlik, Jan Potempa, Tomasz Kantyka, Krzysztof Pyrc

## Abstract

Human coronavirus HKU1 (HCoV-HKU1) is associated with respiratory disease and is prevalent worldwide, but *in vitro* model for virus replication is lacking. Interaction between the coronaviral spike (S) protein and its receptor is the major determinant of virus tissue and host specificity, but virus entry is a complex process requiring a concerted action of multiple cellular elements. Here, we show that KLK13 is required for the infection of the human respiratory epithelium and is sufficient to mediate the entry of HCoV-HKU1 to non-permissive RD cells. We also demonstrated HCoV-HKU1 S protein cleavage by KLK13 in the S1/S2 region, proving that KLK13 is the priming enzyme for this virus. Summarizing, we show for the first time that protease distribution and specificity predetermines the tissue and cell specificity of the virus and may also regulate interspecies transmission. It is also of importance that presented data may be relevant for the emerging coronaviruses, including SARS-CoV-2 and may help to understand the differences in their zoonotic potential.

## INTRODUCTION

Coronaviruses are the largest group within the order *Nidovirales*. Mainly, they cause respiratory and enteric diseases in humans and animals, but some can cause more serious conditions such as hepatitis, peritonitis, or neurological disease. Seven coronaviruses infect humans, four of which (human coronavirus [HCoV]-229E, HCoV-NL63, HCoV-OC43, and HCoV-HKU1) cause relatively mild upper and lower respiratory tract disease and two (SARS-CoV and MERS-CoV) are associated with severe, life-threatening respiratory infections and multiorgan failure (*1–6*). Furthermore, in December 2019 a novel coronavirus SARS-CoV-2 emerged in Hubei province, China, causing pneumonia. To date, almost 90,000 cases were identified and 3,000 patients died worldwide.

Coronaviral infection is initiated by interaction between the trimeric spike (S) protein and its receptor, which is expressed on the surface of the susceptible cell. A number of adhesion and entry receptors have been described for coronaviruses. For example, HCoV-229E (similar to many other alphacoronaviruses) utilizes aminopeptidase N (APN) as the primary entry port (*7*). Surprisingly, its cousin HCoV-NL63 shares receptor specificity with the evolutionarily distant SARS-CoV and SARS-CoV-2: all hijack angiotensin-converting enzyme 2 (ACE2) (*8–11*). HCoV-NL63 was also shown to use heparan sulfate as a primary attachment site (*12–14*). A very different receptor is recognized by MERS-CoV, which binds to dipeptidyl-peptidase 4 (DPP4) (*9, 15, 16*). Another betacoronavirus, HCoV-OC43, binds to N-acetyl-9-O-acetylneuraminic acid (*17, 18*). HCoV-HKU1 remains the great unknown because its cellular receptor has not been identified and all efforts to culture the virus *in vitro* have failed.

HCoV-HKU1 was identified in Hong Kong in 2004. The virus was present in a sample obtained from an elderly patient with severe pneumonia (*19*). Epidemiological studies show a high prevalence of the pathogen in humans; this is because the majority of children seroconvert before the age of 6 years (*20, 21*). While it is not possible to culture HCoV-HKU1 *in vitro,* we and others reported that *ex vivo* fully differentiated human airway epithelium (HAE) and human alveolar type II cells support the infection (*22–25*). A thorough study by Huang X *et al*. demonstrated that HCoV-HKU1 binds to target cells *via O*-acetylated sialic acids on the cell surface; however, this interaction is not sufficient for the infection. The study also showed that the hemagglutinin-esterase (HE) protein of HCoV-HKU1 exhibits sialate-*O*-acetylesterase activity and may act as a receptor-destroying enzyme, thereby facilitating the release of viral progeny (*26*). Bakkers *et al*. proposed that, in order to adapt to the sialoglycome of the human respiratory tract over the evolutionary timescale, HCoV-HKU1 lost the ability to bind to attachment receptors *via* the HE protein (*27*). Recently, Hulswit *et al*. mapped the virus binding site to *O*-acetylated sialic acids, demonstrating that the S1 domain A is responsible for binding to the attachment receptor (*28*).

The S protein is the main player during coronavirus entry, and its characteristics determine the host range. Coronaviral S proteins are class I fusion proteins comprising a large N-terminal ectodomain, a hydrophobic trans-membrane region, and a small C-terminal endodomain. The ectodomain is highly glycosylated and is composed of S1 and S2 domains. The globular S1 domain is highly variable and carries the receptor-binding site, whereas the more conserved rod-like S2 domain undergoes structural rearrangement during entry, which brings the cellular and viral membranes into close proximity. Such a structural switch may be triggered by different stimuli, including receptor binding, proteolytic cleavage of the S protein, and/or a reduction in pH. Because different species require different stimuli, coronaviruses enter cells at different subcellular sites. Some coronaviruses fuse at the plasma membrane, whereas others are believed to enter the cell through receptor-mediated endocytosis, followed by fusion deep within the endosomal compartments (*29–35*). Furthermore, recent reports show that the entry portal may vary depending on tissue/cell characteristics. These differences may affect the host range, pathogenicity, and cell/tissue specificity (*1*).

Host proteases prime coronaviral S proteins. For example, trypsin-mediated cleavage in the small intestine is required for entry of porcine epidemic diarrhea virus (*36*), while a number of coronaviruses from different genera (including HCoV-OC43, HCoV-HKU1, murine hepatitis virus [MHV], MERS-CoV, and infectious bronchitis virus [IBV]) possess a furin cleavage site (*37–41*). Kam *et al.* showed that SARS-CoV S protein can be cleaved by plasmin; however, there is almost no biological evidence for its role *in vivo* (*42*). Cathepsins may also act as S protein-activating enzymes. Indeed, cathepsin L processes the S proteins of SARS-CoV, MERS-CoV, HCoV-229E, and MHV-2 (*43–46*). However, recent reports show that, *in vivo*, respiratory coronaviruses may be activated by the TMPRSS2 protease, which enables endocytosis-independent internalization, thereby re-shaping the entry process (*45, 47–50*). While laboratory strains require priming by cathepsins, S proteins of clinical isolates (e.g., HCoV-229E and HCoV-OC43) undergo TMPRSS2-mediated cleavage at the cell surface, which enables them to fuse with the cell membrane on the cell surface (*51*). A recent study by Shirato *et al*. demonstrates that coronaviruses may lose their ability to infect naturally permissive HAE cultures during cell culture adaptation because the S gene evolves and adjusts to the proteolytic landscape of the immortalized cells (*51*).

Here, we identified a protease belonging to the tissue kallikrein (KLK) family as a new player essential for HCoV-HKU1 entry to the target cell. The KLK family comprises 15 closely related serine proteases with trypsin- or chymotrypsin-like specificity. The expression of these enzymes is tightly regulated, and each tissue has its own unique KLK expression profile. These enzymes play a role in a diverse range of processes during embryonic development to adulthood (*52–56*), and some have been linked to human cancers (*57–60*). The function of some KLKs remains to be elucidated, but obtained results suggest that protease distribution may be an important factor pre-determining the cell and tissue specificity of the virus, but also regulating the interspecies transfers.

Further, the data presented herein bring us a step closer to developing a convenient *in vitro* culture model and possibly identifying the cellular receptor for this virus. It is also of importance that presented data may be relevant for the emerging coronaviruses and may help to understand the differences in their zoonotic potential.

## RESULTS

### Several KLKs are upregulated after infection of HAE with HCoV-HKU1

First, we asked whether HCoV-HKU1 infection modulates the expression of human KLKs. HAE cultures were infected with HCoV-HKU1 or mock-inoculated. At 120 h post-inoculation (p.i.), cells were collected and the expression of mRNAs encoding KLKs was analyzed. We detected the expression of KLK7, KLK8, KLK10, KLK11, and KLK13 in non-infected fully differentiated cultures. However, the pattern in HCoV-HKU1-infected cells was different: we detected an upregulation of KLK7, KLK8, KLK10, KLK11 and KLK13. Further, KLK1, KLK5, KLK6, KLK9, KLK12 and KLK14 were expressed in the infected cells. KLK2, KLK3, and KLK15 were not expressed (**Fig. 1**).

**Figure 1.**
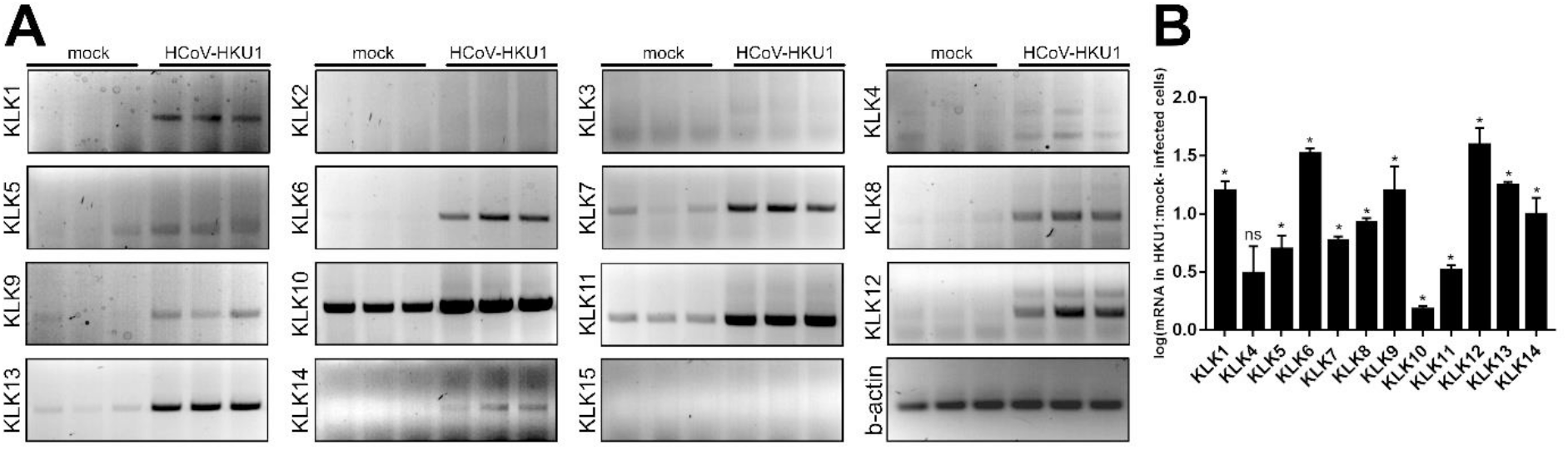
Up-regulation of several KLKs after HCoV-HKU1 infection in HAE. HAE cultures were infected with HCoV-HKU1 (10^6^ RNA copies per ml) or mock for 2 h at 32°C and cultured for 5 days. Cellular RNA was isolated, treated with DNase, reverse transcribed and each KLKs mRNA was amplified using specific primers. The analysis was performed two times using cells obtained from different donors, each time in triplicate. (**A**) Amplified PCR products were resolved and detected in 1.5% (w:v) agarose gel in 1 × TAE buffer. (**B**) The expression of each KLK comparing to β-actin control was assessed semi-quantitively by densitometry and is presented as a log change of signal specific for KLKs mRNA in HCoV-HKU1-infected cells, compared to the mock-infected cells. The experiment was performed three times using cells from different donors, each time with three biological replicates. For comparisons by Student’s t-test, *Indicates P < 0.05; ns, not significant.

### KLK13 is essential for HCoV-HKU1 infection

S protein priming is a prerequisite for coronavirus entry; therefore, we tested whether KLKs take part in this process by culturing cells in the presence/absence of KLK inhibitors (**Table 1**) (*61*). For this, HAE cultures were pre-incubated with each inhibitor (10 µM) or with vehicle control (DMSO) and mock-inoculated or inoculated with the virus (10^6^ RNA copies per ml) in the presence of the inhibitor. Apical washes were collected each day for analysis of virus replication. Subsequently, viral RNA was isolated and reverse transcribed (RT), and the HCoV-HKU1 yield was determined by quantitative real-time PCR (qPCR). The results showed that HCoV-HKU1 replication was inhibited in the presence of a KLK13 inhibitor; this was not the case for cells treated with DMSO or with inhibitors specific for KLK7 or KLK8 (**Fig. 2A**).

**Figure 2.**
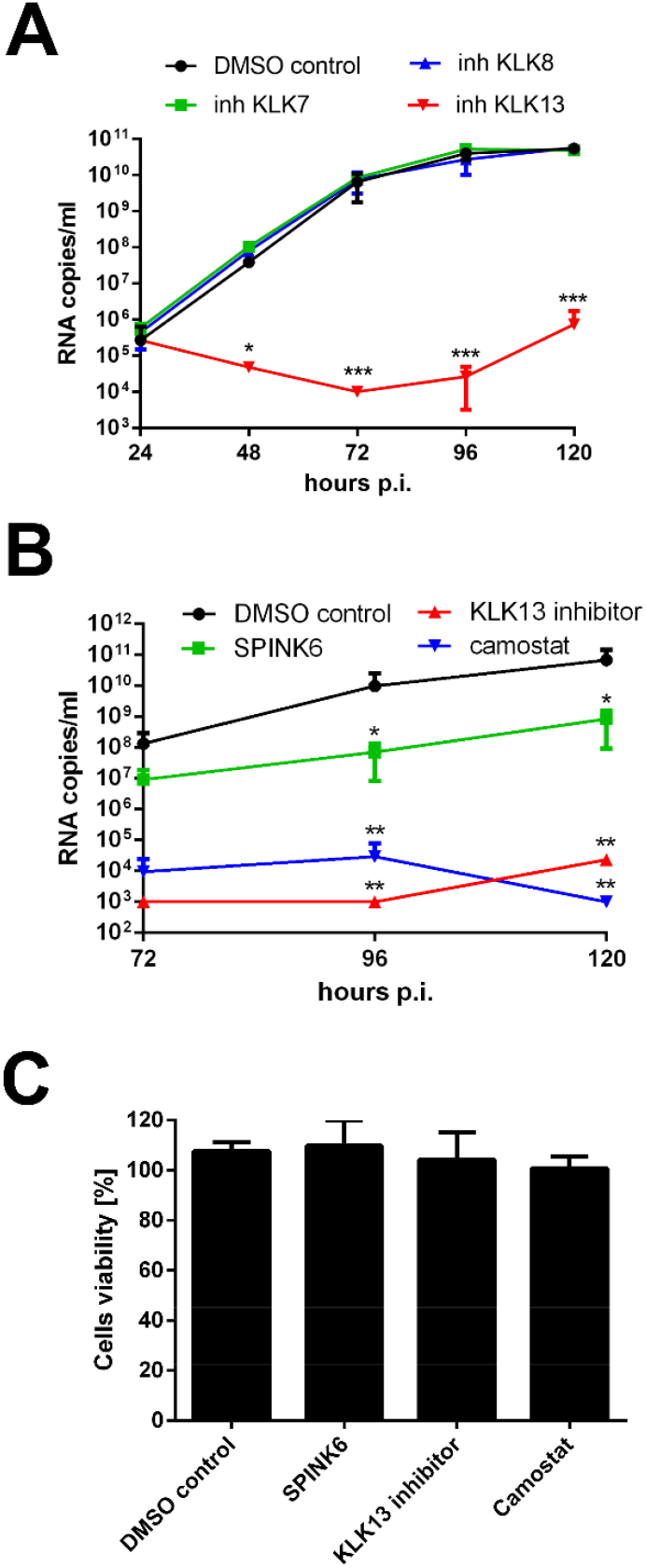
HCoV-HKU1 infection is dependent on KLK13 activity. HAE cultures were inoculated with HCoV-HKU1 (10^6^ RNA copies per ml) for 2 h at 32°C in the presence of 10 μM KLKs inhibitors (**Table 1**) or DMSO (**A**); 10 μg/ml SPINK6, 10 μM KLK13 inhibitor, 100 μM camostat or DMSO (**B**). To analyze virus replication kinetics, each day post-infection, 100 μl of 1 × PBS with a given inhibitor was applied to the apical surface of HAE cultures and collected after 10 min incubation at 32°C. Replication of HCoV-HKU1 was evaluated using an RT-qPCR and the data are presented as RNA copy number per ml. The assay was performed twice, each time in triplicate, and average values with standard errors are presented. Statistical significance was assessed with the Student’s t-test, and the asterisks indicate: * P < 0.05, ** P < 0.005, *** P < 0.0005. (**C**) Cytotoxicity of inhibitors in HAE cultures. Cell viability was assessed with the XTT assay on mock-treated cells at 120 h post-infection. Data on the y-axis represent the percentage values obtained for the untreated reference samples. The assay was performed in triplicate and average values with standard errors are presented.

**Table 1.** KLKs inhibitors used in the study (61).

| Protein | Inhibitor sequences |
| --- | --- |
| **KLK7** | Biotin-KTLF-CMK / Biotin-PEG-KTLF-CMK |
| **KLK8** | Biotin-TNKR-CMK / Biotin-PEG-TNKR-CMK |
| **KLK13** | Biotin-VRFR-CMK / Biotin-PEG-VRFR-CMK |

Next, we analyzed HCoV-HKU1 replication in the presence of a family-specific KLK inhibitor SPINK6 at a concentration of 10 µg/ml (*62, 63*) or 100 µM camostat (a broad inhibitor of serine proteases, also a potent inhibitor of KLK13) (*34, 64*). We noted inhibition of

HCoV-HKU1 replication in the presence of both inhibitors (**Fig. 2B**). All inhibitors were used at non-toxic concentrations (**Fig. 2C**).

The experiments conducted so far suggested that KLK13 is required for virus infection. However, one may question the specificity of the KLK13 protease inhibitors. To ensure that KLK13 is indeed the priming enzyme during HCoV-HKU1 infection, we developed HAE cultures by transforming cells with lentiviral vectors encoding shRNAs targeting KLK13 mRNA. We then confirmed that the expression of the protease was silenced (HAE_shKLK13). Non-modified HAE cultures (HAE_ctrl), cultures modified using a lentiviral vector to express the GFP protein (HAE_GFP), and HAE cultures transduced with an empty lentiviral vector (HAE_vector) were used as controls. Following transduction and differentiation, expression of KLK13 mRNA in HAE_shKLK13 was almost undetectable, in contrast to the control cultures (**Fig. 3A**). Importantly, HAE_shKLK13 cells continued to differentiate and formed pseudostratified cultures (**Fig. 3B**). Next, we infected HAE_ctrl, HAE_GFP, HAE_vector, and HAE_shKLK13 with HCoV-HKU1 (10^6^ RNA copies per ml) and incubated them for 2 h at 32°C with the viral stock solution. Cultures were maintained at 32°C for 5 days at an air-liquid interface. Apical washes were collected, and virus yield was determined by RT-qPCR. We found that, in contrast to that in control cultures, replication of virus in HAE_shKLK13 was abolished (**Fig. 3C**). Overall, these data suggest that silencing the *KLK13* gene in HAE inhibits virus infection, indicating that KLK13 is necessary for HCoV-HKU1 infection.

**Figure 3.**
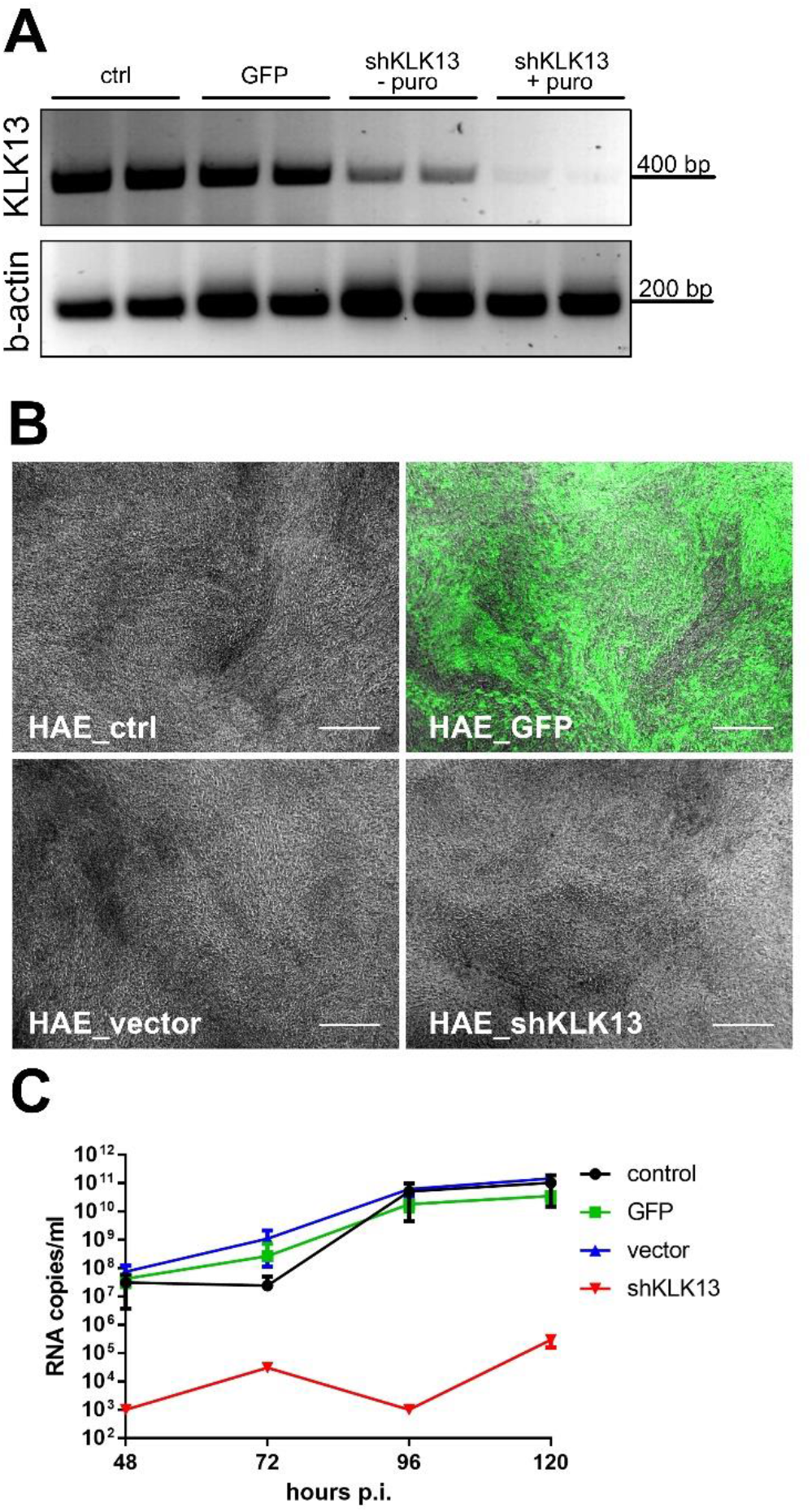
HCoV-HKU1 does not replicate in HAE cells deficient in KLK13. Primary human epithelial cells were transduced with lentiviral vectors harboring the GFP (HAE_GFP) protein, empty pLKO.1-TRC vector (HAE_vector) or shRNA for KLK13 mRNA (HAE_shKLK13). As a control, not transduced HAE cultures were used (HAE_ctrl). (**A**) The KLK13 mRNA was evaluated before (- puro) and after puromycin (+ puro) selection of positively transduced cells, β-actin was used as an internal control. (**B**) Microscopic examination of all HAE cultures after 4 weeks culture in ALI at 37°C. Scale bar = 200 μm. (**C**) All HAE cultures were inoculated with HCoV-HKU1 (10^6^ RNA copies per ml) for 2 h at 32°C and cultured for 5 days. Each day post-infection, 100 μl of 1 × PBS was applied to the apical surface of HAE cultures and collected after 10 min incubation at 32°C. Replication of HCoV-HKU1 was evaluated using an RT-qPCR and the data are presented as RNA copy number per ml. The assay was performed twice, each time in triplicate, and average values with standard errors are presented.

### KLK13 enables entry of HCoV-HKU1 pseudoviruses

We determined that KLK13 is essential for efficient HCoV-HKU1 infection in HAE cultures and we started to wonder whether this enzyme may be a determinant of the cell and tissue specificity of the virus. Previous studies showed that RD cells support virus attachment *via* sialic acids, but this does not allow for the virus entry (*26*). To test whether cell surface proteases my render RD cells permissive, we generated RD cells expressing human KLK13 or TMPRSS2 proteases. RD cells were transduced with lentiviral vectors harboring the KLK13 gene (RD_KLK13), control vector (RD_ctrl), or TMPRSS2 (RD_TMPRSS2). Due to the lack of KLK13 specific antibodies, we verified its presence based on RT-PCR (**Fig. 4A**). The presence of TMPRSS2 in RD_TMPRSS2 cells was confirmed using Western blot. The TMPRSS2 band in RD cells was observed at 25 kDa, which corresponds to one of the naturally occurring splicing variants (**Fig. 4B**). Subsequently, we transduced RD_ctrl, RD_KLK13 and RD_TMPRSS2 cells with HIV particles pseudotyped with HCoV-HKU1 S glycoprotein (S-HKU1), control VSV G protein (VSV-G) or lacking the fusion protein (ΔEnv). After 3 day culture at 37°C, pseudovirus entry was quantified by measurement of the luciferase activity. As shown in **Fig. 4C**, all cultures were effectively transduced with control VSV-G vectors, while only RD_KLK13 cells were permissive to S-HKU1 pseudoviruses. This clearly showed that KLK13, and not TMPRSS2 is involved in HCoV-HKU1 entry. Furthermore, S-HKU1,d VSV-G and ΔEnv pseudoviruses were overlaid onto fully differentiated HAE cultures in the presence of KLK13 inhibitor (10 µM) or DMSO. After 3 day culture at 37°C pseudovirus entry was quantified by measurement of the luciferase activity. Despite low transduction efficiency in HAE, we observed an increase in luciferase activity in cultures treated with S-HKU1 pseudoviruses, compared to ΔEnv, which was completely abolished in the presence of KLK13 inhibitor (**Fig 4D**). Overall, these data demonstrated that KLK13 activity drives HCoV-HKU1 entry into cells.

**Figure 4.**
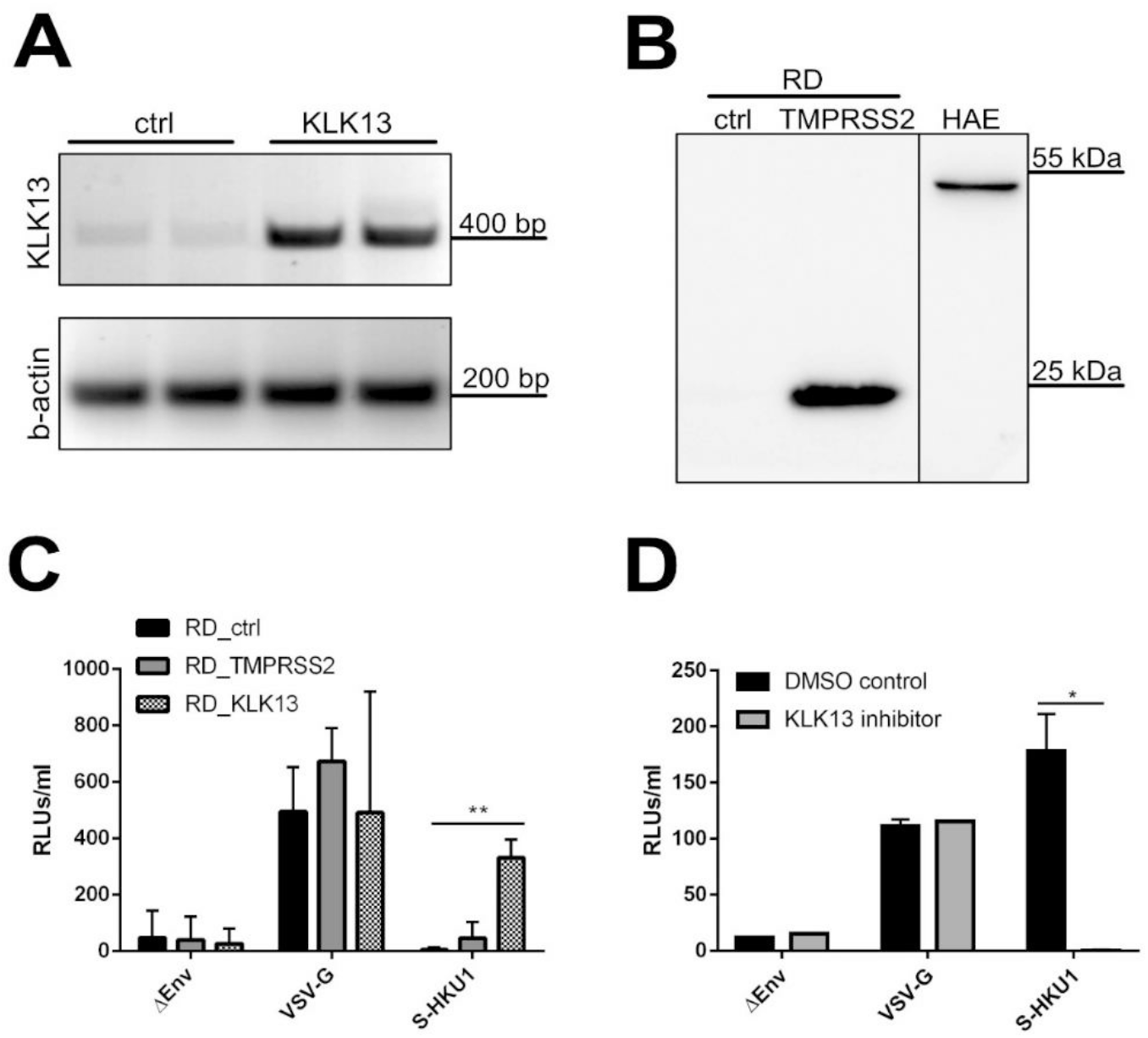
RD cells expressing KLK13 are permissive for the HCoV-HKU1 pseudoviruses. (**A**) RD cells were transduced with lentiviral vectors harboring the KLK13 gene (KLK13) or empty vector (ctrl). The presence of KLK13 mRNA was evaluated in RD cells after blasticidin selection; β-actin was used as an internal control. (**B**) RD cells were transduced with lentiviral vectors harboring the TMPRSS2 gene (TMPRSS2) or empty vector (ctrl). After blasticidin selection, cells were lysed and proteins were analyzed with SDS-PAGE. TMPRSS2 was detected in RD cell lysates (50 μg of protein per lane) and HAE cultures lysate (25 μg of protein per lane) using the specific antibody. (**C**) RD control (RD_ctrl), TMPRSS2-expressing (RD_TMPRSS2) or KLK13-expressing (RD_KLK13) cells were transduced with HIV pseudoviruses decorated with VSV-G protein (VSV-G), S-HKU1 glycoprotein (S-HKU1) or control viruses without the fusion protein (ΔEnv). After 72 h at 37°C, the entry of pseudoviruses was measured by means of luminescence signal in cell lysates. The assay was performed twice, each time in triplicate, and average values with standard errors are presented. For comparisons by Student’s t-test, **Indicates P < 0.005. (**D**) HAE cultures were inoculated with HIV pseudoviruses harboring VSV-G control protein, S-HKU1 or control viruses without the fusion protein (ΔEnv) in the presence of KLK13 inhibitor (10 μM) or DMSO. After 72 h at 37°C the entry of pseudoviruses was measured by means of luminescence signal in cell lysates (RLUs per ml of lysate sample). The assay was performed in duplicate, and average values with standard errors are presented. Statistical significance was assessed with the Student’s t-test, and the asterisk indicates P < 0.05.

### KLK13 enables the replication of HCoV-HKU1 in RD cells

Obtained results showed that KLK13 expression on RD cells was sufficient for HCoV-HKU1 pseudovirus entry. Here, we aimed to test whether KLK13 presence renders RD cells permissive for HCoV-HKU1 infection. For this, we infected RD_ctrl and RD_KLK13 cells with HCoV-HKU1 (10^8^ RNA copies per ml) and incubated the culture for 7 days at 32°C in the presence or absence of a KLK13 inhibitor (10 µM) or DMSO. Next, cellular RNA was isolated and the presence of HCoV-HKU1 N subgenomic mRNA (N sg mRNA), which is considered to be a hallmark of coronaviral infection, was assessed (*14*). sg mRNA appeared in RD_KLK13 cells, while no signal was detected in cultures supplemented with the KLK13 inhibitor nor in RD_ctrl cells (**Fig. 5A**).

**Figure 5.**
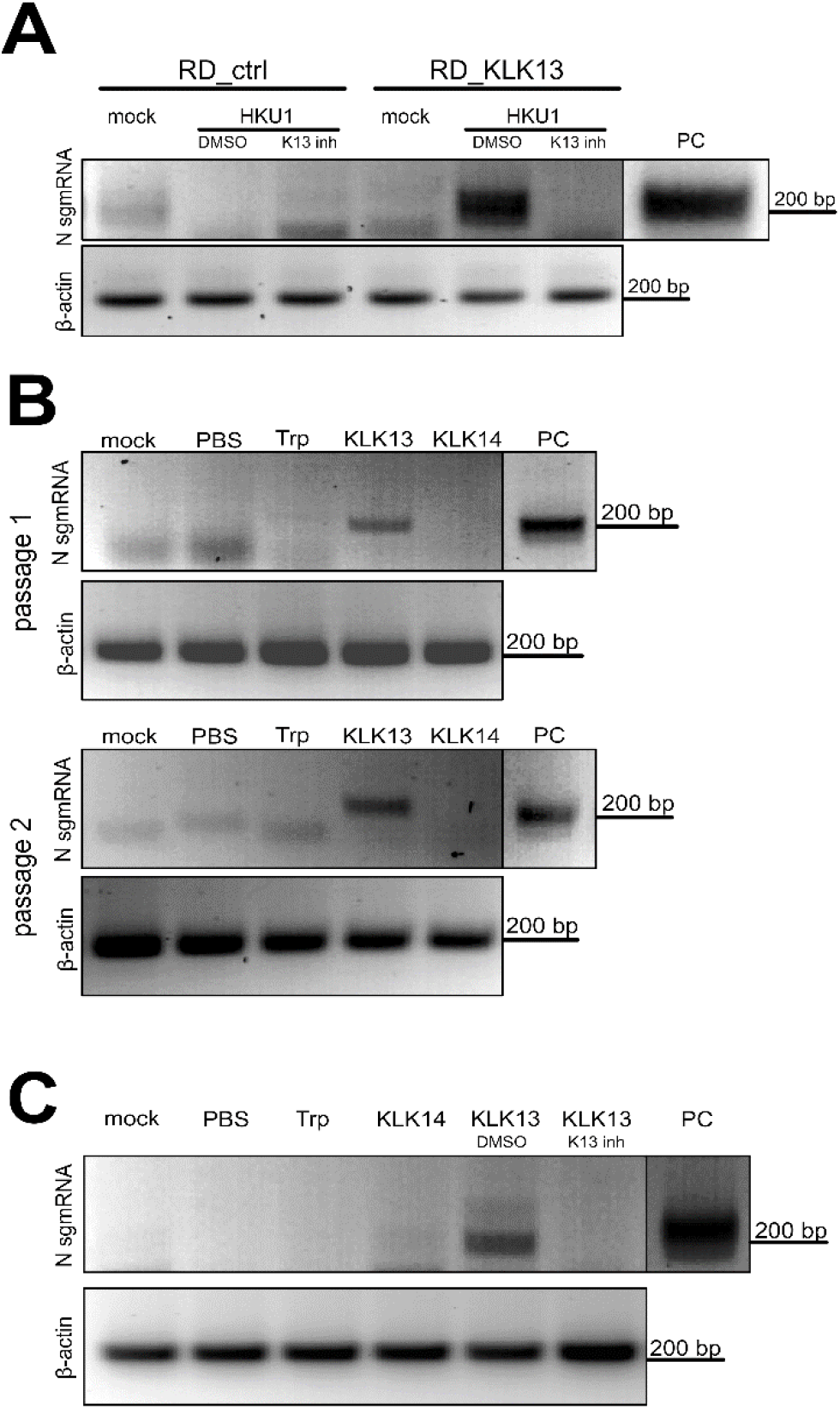
HCoV-HKU1 replicates in RD cells expressing KLK13 protease. (**A**) Control (RD_ctrl) or KLK13-expressing (RD_KLK13) cells were inoculated with HCoV-HKU1 (10^6^ RNA copies per ml) or mock in the presence of 10 μM KLK 13 inhibitor (K13 inh) or control DMSO. After 7 days culture at 32°C, total RNA was isolated, reverse transcribed and subgenomic mRNA for N protein was detected in semi-nested PCR, β-actin was used as an internal control. PC = positive control from virus-infected HAE cells. (**B**) HCoV-HKU1 was incubated with 200 nM trypsin (Trp), KLK13, KLK14 or PBS for 2 h at 32°C and further applied on the RD cells. After 7 days at 32°C total RNA was isolated, reverse transcribed and subgenomic mRNA for N protein was detected in semi-nested PCR (passage 1). Simultaneously, 1 ml of cell culture supernatant was harvested and applied to freshly seeded RD cells with medium supplemented with fresh enzymes. After 7 days at 32°C, subgenomic mRNA for the N protein was detected in semi-nested PCR (passage 2). β-actin was used as an internal control. (**C**) HCoV-HKU1 was incubated with 200 nM trypsin (Trp), KLK14, KLK13 with KLK13 inhibitor (K13 inh), control DMSO (DMSO), or PBS for 2 h at 32°C and further applied onto the RD cells. Subgenomic mRNA for N protein was detected in semi-nested PCR; β-actin was used as an internal control. PC: positive control from virus-infected HAE cultures.

To further confirm the role of KLK13 during HCoV-HKU1 infection RD cells were supplemented with purified human KLK13 or KLK14 (*61*). The latter was used as a negative control. Virus stock was incubated for 2 h at 32°C with 200 nM KLK13, KLK14, or trypsin or control (PBS). Next, RD cells were incubated for 7 days at 32°C with the virus (diluted 10-fold in DMEM) or mock samples (in DMEM), after which cellular RNA was isolated and HCoV-HKU1 infection was analyzed by means of N sg mRNA detection. Again, we found that N sg mRNA was produced only in the presence of KLK13 (**Fig. 5B**). Further, we passaged HCoV-HKU1 twice in RD cells. Briefly, 1 ml of cell culture supernatant from the first experiment was transferred to fresh RD cells and fresh enzymes were added (final concentration 200 nM). Cultures were then incubated at 32°C for 7 days. Cellular RNA was isolated and HCoV-HKU1 infection was monitored by detecting N sg mRNA. The infection occurred only in the presence of KLK13 (**Fig. 5B**). However, we observed no cytopathic effects (CPEs), and replication levels were very low (no significant increase over control levels on RT-qPCR; data not shown). To further test the effect of KLK13 on replication of HCoV-HKU1 in RD cells, the virus stock was incubated with purified KLK13 (200 nM) and incubated in the presence or absence of a KLK13 inhibitor (10 µM) or DMSO. After 2 h at 32°C pre-treated virus stock was diluted in media as described above and overlaid on RD cells. After 7 days at 32°C we evaluated the presence of the HCoV-HKU1 N sg mRNA. The virus replicated only after treatment with KLK13, and supplementation with the inhibitor blocked this effect infection (**Fig. 5C**).

### KLK13 primes the HCoV-HKU1 S protein

Expression of KLK13 by cells previously resistant to HCoV-HKU1 renders them susceptible; therefore, we asked whether this is due to proteolytic activation of the S protein. We tested this using the CleavEx method, in which a peptide of interest is exposed in the N-terminal region of the proteolytically-resistant HmuY carrier protein. Briefly, two-hybrid His-tagged CleavEx proteins were prepared, both harboring 8-amino acid peptide sequences of the HCoV-HKU1 S protein. The first peptide contained the S1/S2 cleavage site (amino acids 757–764), which in some coronaviruses is activated during protein biosynthesis, during virus exocytosis, or after receptor engagement. The second harbored the S2/S2’ cleavage site (amino acids 901–908), which is an additional region prone to proteolytic cleavage (*25, 38*). Proteins were purified and further incubated for 3 h at 37°C with increasing concentrations of purified KLK13. Subsequently, proteins were resolved by SDS-PAGE and detected by western blotting with antibodies specific for His-tagged proteins. The analysis showed that, in the presence of 500 nM KLK13, the CleavEx protein harboring the S1/S2 cleavage site was degraded; however, the CleavEx protein harboring the S2/S2’ site remained intact. Because KLKs are produced as pro-forms that undergo self-activation, an additional band of His-tagged purified pro-KLK13 (HisTag-pro-KLK13) was observed after treatment with 500 nM KLK13 (**Fig. 6A**). The product of the S1/S2 cleavage was further sequenced by N-terminal Edman degradation showing the following sequence: R↓SISA, which corresponds to the S1/S2 site. This result shows clearly that the S1/S2 region of the HCoV-HKU1 S protein is prone to KLK13-mediated cleavage.

**Figure 6.**
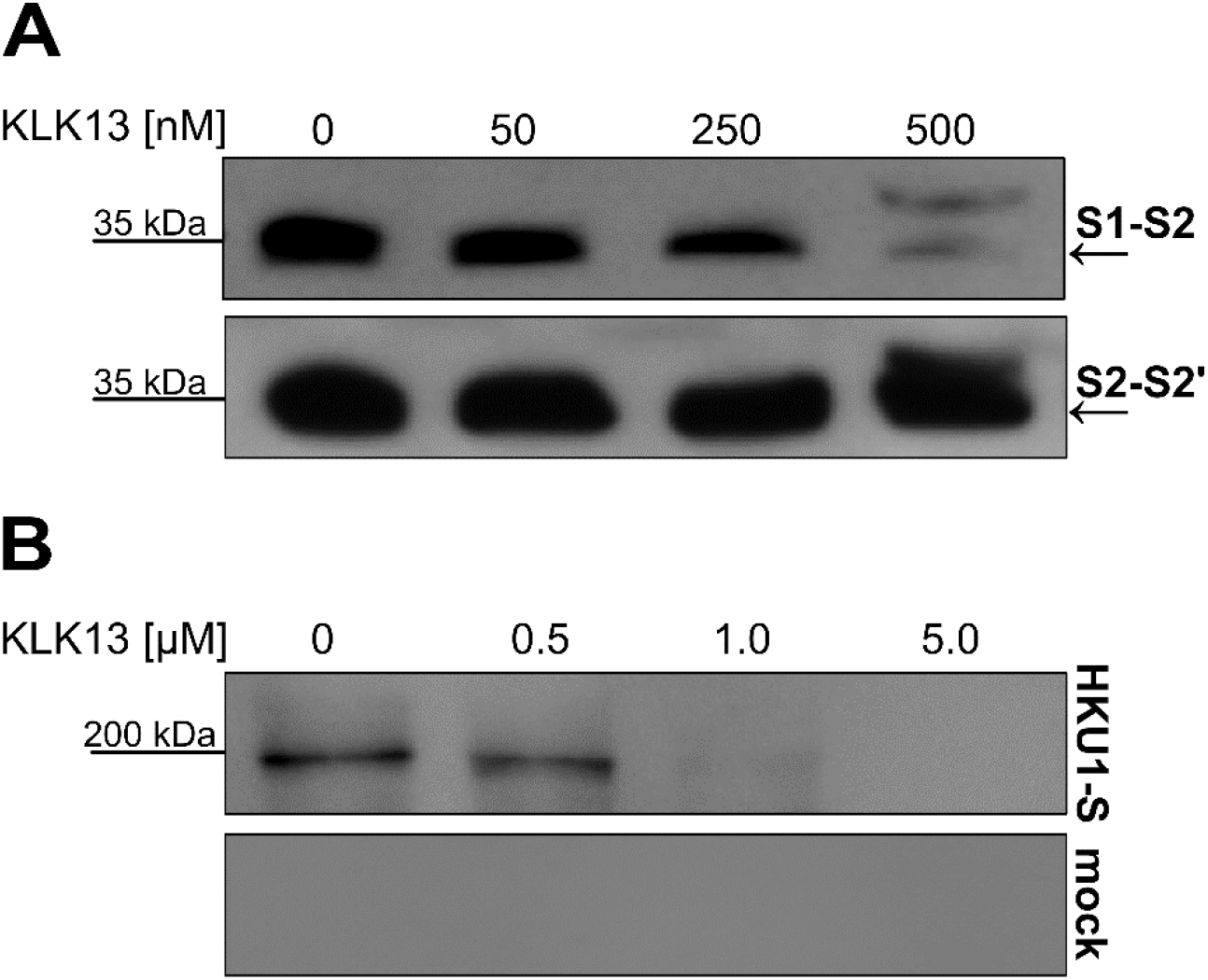
KLK13 cleaves the HCoV-HKU1 Spike protein between S1 and S2 domains. (**A**) 15 ng of CleavEx proteins harboring the S1/S2 or S2/S2’ site were incubated at 37°C for 3 h with different concentrations of the purified KLK13. (**B**) The full-length HKU1-S protein or mock was incubated at 37°C for 3 h with different concentrations of the purified KLK13. After samples denaturation at 95°C, proteins were resolved by SDS-PAGE and detected using the horseradish peroxidase-labeled anti-His-tag antibody.

Furthermore, we aimed to confirm the cleavage using a full-length Spike protein of HCoV-HKU1 (HKU1-S). For this, we expressed the HKU1-S in 293T cells, purified the protein using 6 × His tag, and incubated for 3 h at 37°C with increasing concentrations of purified KLK13. Subsequently, HKU-S or mock proteins were resolved by SDS-PAGE and detected by western blotting with antibodies specific to the tag. The analysis showed that in the presence of 1 µM KLK13 the HKU1-S was degraded (**Figure 6B**). The S protein was observed at ∼150 kDa, which is consistent with the migration speed reported for these highly glycosylated proteins (*1*).

## DISCUSSION

Receptor recognition is the first, essential step of the virus infection process. The coronaviral S protein mediates virus entry into host cells by binding to a specific receptor. A combination of stimuli, e.g., receptor binding, proteolytic cleavage, and exposure to low pH results in rearrangement of the S protein and, consequently, to membrane fusion and virus entry (*1*). Although the structure of both the HCoV-HKU1 S ectodomain and the receptor-binding domain has been resolved, the receptor determinant remains unknown (*28, 38, 65*). In a previous study, we showed that HCoV-HKU1 utilizes *O*-acetylated sialic acids on host cells as an attachment receptor (*26*). Here, we present data demonstrating that the protease KLK13 is required for HCoV-HKU1 infection of the respiratory epithelium.

Human KLKs take part in multiple physiological processes, including skin desquamation, tooth enamel formation, kidney and brain function, and synaptic neural plasticity (*66–75*). Also, recent studies demonstrate a role for some KLKs during viral infections. For instance, KLK8 plays a role in the proteolytic activation of the human papillomavirus capsid protein, thereby mediating virus entry into host cells (*76*). Also, KLK5 and KLK12 are secreted into the respiratory tract, where they support replication of the influenza A virus by cleaving the hemagglutinin protein (*77, 78*); however, these proteins belong to a large pool of cell surface proteases, the orchestrated action of which promotes virus replication.

Here, we found that the yield of HCoV-HKU1 from HAE fell in the presence of SPINK6 (inhibitor of KLK13) (*62, 63*) and camostat (a broad range inhibitor of serine proteases). However, the first compound also inhibits other KLKs (*63*), while the second block the activity of a wide range of serine proteases and was used previously to demonstrate the role of TMPRSS2/4 proteases during viral infection (34, 45, 51, 64, 79, 80). The relatively low inhibition of HCoV-HKU1 replication in the presence of SPINK6 possibly results from non-optimal compound concentration at the HAE cultures, and cytotoxicity at higher concentrations(*63*). Broad spectrum protease inhibitors are now used widely for virus research, although their non-specific activity makes the results equivocal. For example, Matsuyama *et al*. showed recently that the furin inhibitor dec-RVKR-CMK interferes with the activity of several proteases, and that its previously described inhibitory activity during MERS-CoV infection is not specific to furin; instead, its activity is due to non-specific inhibition of cathepsin L and TMPRSS2 (*81*). We tried to use specific KLK inhibitors developed in our laboratory (*61*). Considering the small arsenal of tools available to researchers studying KLKs, only three compounds were readily available. Treating HAE cultures with these inhibitors revealed that only compounds designed to inhibit KLK13 hampered HCoV-HKU1 replication. However, the great similarity between different KLKs makes one doubt the specificity of these inhibitors, despite their performance in biochemical assays. Therefore, we decided to silence KLK13 in HAE cultures. This abolished virus replication *ex vivo*, thereby confirming the importance of KLK13 during infection. KLK13 is thought to be secreted and membrane-bound(*82, 83*).

This study showed that HCoV-HKU1 infection in HAE modulates the expression of different KLKs, including KLK13. In our study, KLKs expression was tested using semi-quantitative PCR, and for that reason, we were unable to show the level of KLKs modulation after HCoV-HKU1. However, the pattern of virus-induced expression of several KLKs could be observed. KLK mechanism of activation is a complex process and until now it has only been proven that most KLK genes are regulated by steroids and other hormones (*84*). It is also important to remember, that KLK expression is regulated in a similar manner, and the induction of a single gene usually results in overexpression of the whole cluster (*85, 86*). While one may assume that the virus stimulates KLK13 production to promote the infection, this up-regulation is likely a natural response of the damaged tissue, as KLKs were previously reported to take part also in tissue regeneration (*87–89*). Further, increased expression of KLKs may be the response to the inflammatory process, as Seliga *et al* demonstrated that KLK-kinin system is a potent modulator of innate immune responses (*90*).

The experiments performed herein show the importance of KLK13 for virus entry into susceptible cells; therefore, we speculated that scattered distribution of different KLKs in different tissues may be one of the determinants of the HCoV-HKU1 tropism (*53, 91*). We tested the purified enzyme expressed in the eukaryotic cells; however, we also developed a cell line constitutively expressing the enzyme. As an *in vitro* model for our studies, we used RD cells previously reported to carry attachment receptors for the virus (*26*). Here, using pseudoviruses decorated with S-HKU1 proteins we showed that KLK13 presence on RD cells is sufficient for virus entry and renders these cells permissive. Our experiments also showed that in contrast to previous reports, TMPRSS2 is not involved in this process(*51*). Furthermore, we observed that RD cells supported the replication of the virus in the presence of KLK13 and that this effect was reversed in the presence of the specific KLK13 inhibitor. We were, however, not able to culture the virus to high yields. HCoV-HKU1 replication in KLK13-expressing RD cells remained inefficient and RTqPCR assessment did not reveal significant increases in the amounts of viral RNA. For that reason, we are only able to detect viral sg mRNAs, which are considered to be the hallmark of coronaviral replication.We believe that this may be due to non-optimal infection conditions, which may include inappropriate KLK13 concentrations or low density of the entry receptor. Also, it is possible that RD cells may not support efficient replication of the virus due to factors unrelated to the entry process. Nonetheless, our results show that the HCoV-HKU1 entry receptor is present on RD cells, and we were able to trigger virus entry and replication; these findings warrant further exploration.

Most coronaviral S proteins are processed into S1 and S2 subunits by host proteases, which allows conformational changes in the S protein and leads to fusion of the viral and cellular membranes (*1, 92*). As shown in the recent work by Kirchdoerfer *et al*., the HCoV-HKU1 S protein has two regions that are prone to proteolytic activation: the S1/S2 furin cleavage site and a secondary cleavage site termed S2’, which is adjacent to a potential fusion peptide (*38*). While the S1/S2 site is believed to be processed by furin during protein biosynthesis, the S2/S2’ site is expected to be cleaved during virus entry. As we already knew that KLK13 is sufficient for HCoV-HKU1 infection of naturally non-permissive RD cells, we aimed to investigate whether this was the direct result of KLK13-mediated proteolytic cleavage of the S protein. For this, we employed the CleavEx method, in which peptide of interest is coupled to the carrier HmuY protein and then undergoes proteolytic cleavage by the enzyme being tested. We found that the S1/S2 site was efficiently cleaved by KLK13, whereas the S2/S2’ region remained intact. As CleavEx technique is a convenient surrogate system allowing for precise mapping of the cleavage site, it has some limitations. To ensure the reliability of results, purified full-length HCoV-HKU1 S protein was subjected to the proteolytic cleavage. Also here we observed efficient cleavage of the HCoV-HKU1 S protein by KLK13.

While the results presented here show that KLK13 is able to process the HCoV-HKU1 S protein, one may question whether the cleavage is sufficient for HCoV-HKU1 entry. It was previously presented for MERS-CoV that two consecutive enzymatic scissions are required for activation of the S protein. In this scenario, KLK13 would prime the HCoV-HKU1 S at S1-S2 site, enabling scission at S2-S2’ site by the TMPRSS2 or another host protease (*38, 40, 93*). This may be one of the factors limiting the HCoV-HKU1 replication in RD_KLK13 cells, as only minimal replication is observable.

Summarizing, we show that KLK13 is a key determinant of HCoV-HKU1 tropism. This may explain why, since its first identification in 2004, all efforts to culture HCoV-HKU1 in standard cell lines have failed. We believe that this study increases our knowledge of HCoV-HKU1 and may promote the future in-depth investigation of coronaviruses. Considering the increasing number and diversity of coronaviruses, and the proven propensity of coronaviruses to cross the species barrier and cause severe diseases in humans, further research on the role of different proteases in coronaviral infections is necessary.

## MATERIALS AND METHODS

### Plasmid constructs

KLK13 and TMPRSS2 genes were amplified by PCR using cDNA obtained from HAE cells. Each PCR product was cloned into pWPI plasmid for lentivirus production and sequence verified. pLKO.1-TRC cloning vector was a gift from David Root (Addgene plasmid # 10878)(*94*). Oligonucleotides for the generation of shRNA against KLK13 (3 different shRNAs targeting the exons encoding the active site) were hybridized and cloned into pLKO.1-TRC vector. The full-length HKU1-S gene was amplified by PCR using pCAGGS/HKU1-S plasmid that was a gift from Xingchuan Huang. The PCR product was cloned into pSecTag2 cloning vector and sequence verified. Primer sequences are provided in **Table 2**.

**Table 2.**
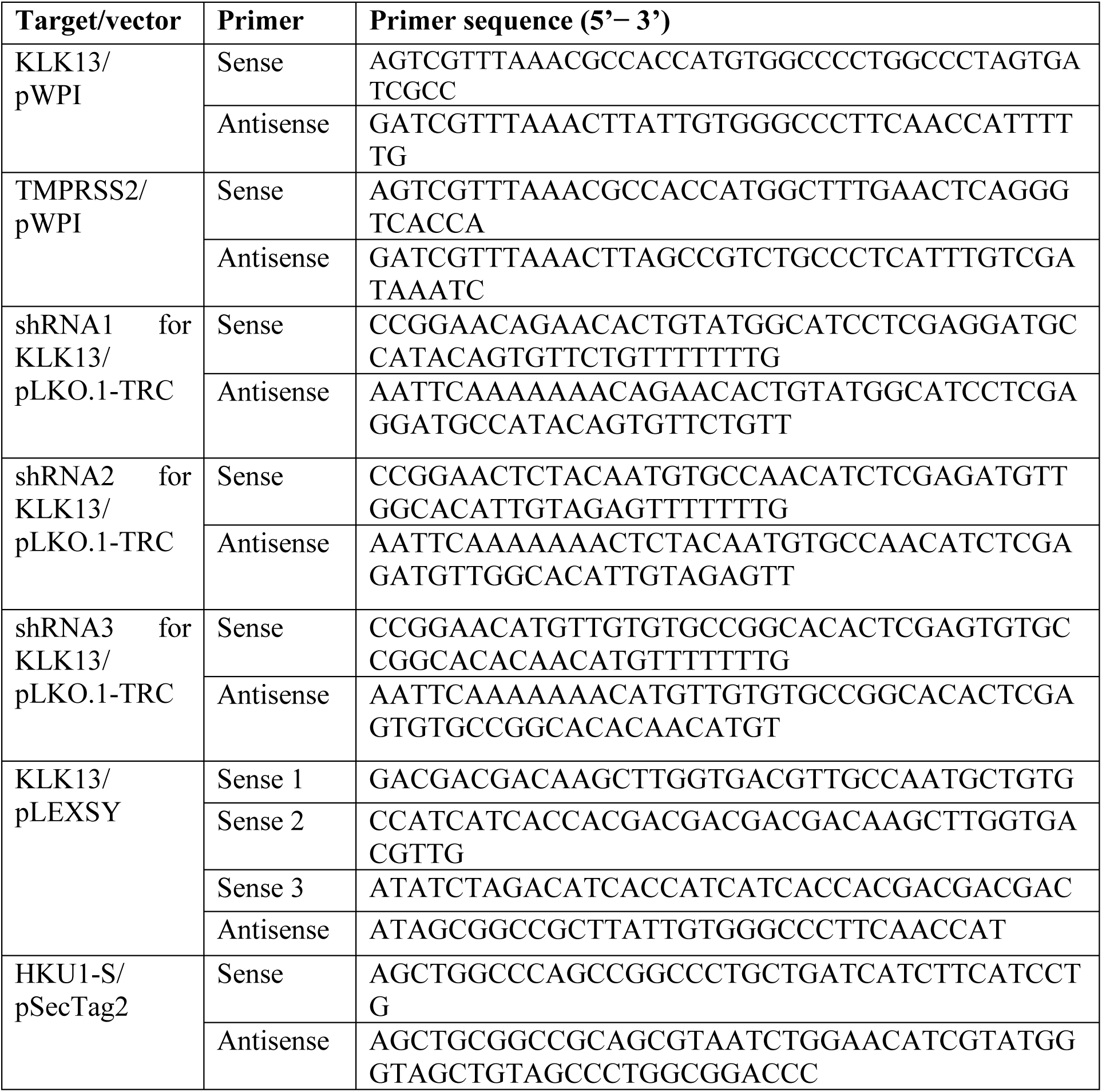
Primers used for generation of plasmid constructs.

### Cell culture

RD (*Homo sapiens* muscle rhabdomyosarcoma; ATCC: CCL-135) and 293T (*Homo sapiens* kidney epithelial; ATCC: CRL-3216) cells were cultured in Dulbecco’s MEM (Thermo Fisher Scientific, Poland) supplemented with 3% fetal bovine serum (heat-inactivated; Thermo Fisher Scientific, Poland) and antibiotics: penicillin (100 U/ml), streptomycin (100 μg/ml), and ciprofloxacin (5 μg/ml). Cells were maintained at 37°C under 5% CO_2_.

### Human airway epithelium (HAE) cultures

Human epithelial cells were isolated from conductive airways resected from transplant patients. The study was approved by the Bioethical Committee of the Medical University of Silesia in Katowice, Poland (approval no: KNW/0022/KB1/17/10 dated 16.02.2010). Written consent was obtained from all patients. Cells were dislodged by protease treatment, and later mechanically detached from the connective tissue. Resulting primary cells were first cultured in the selective media to proliferate in the presence of the Rho-associated protein kinase (ROCK) inhibitor (Y-27632, 10 µg/ml, Sigma-Aldrich, Poland) (*95*). Further, cells were trypsinized and transferred onto permeable Transwell insert supports (ϕ = 6.5 mm). Cell differentiation was stimulated by the media additives and removal of media from the apical side after the cells reached confluence. Cells were cultured for 4-6 weeks to form well-differentiated, pseudostratified mucociliary epithelium (*96*). All experiments were performed in accordance with relevant guidelines and regulations.

### Cell viability assay

HAE cultures were prepared as described above. Cell viability assay was performed by using the XTT Cell Viability Assay (Biological Industries, Israel), according to the manufacturer’s instructions. Briefly, on the day of the assay 100 μl of the 1 × PBS with the 30 μl of the activated XTT solution was added to each well/culture insert. Following 2 h incubation at 37°C, the solution was transferred onto a 96-well plate and the signal was measured at λ = 490 nm using the colorimeter (Spectra MAX 250, Molecular Devices). The obtained results were further normalized to the control sample, where cell viability was set to 100%.

### Virus infection

HAE cultures were washed thrice with 100 μl of 1 × PBS, following inoculation with HCoV-HKU1 (strain Caen 1) or mock (cell lysate). After 2 h incubation at 32°C unbound virions were removed by washing with 100 μl of 1 × PBS and HAE cultures were cultured at air-liquid interphase until the end of the experiment. Due to the lack of a permissive cell line it was not possible to titrate the virus stock for infection experiments and therefore the inoculum was quantified using RT-qPCR.

RD cells grown in 90% confluency were infected with HCoV-HKU1 (10^8^ RNA copies per ml) in Dulbecco’s MEM (Thermo Fisher Scientific, Poland) supplemented with 3% fetal bovine serum (heat-inactivated; Thermo Fisher Scientific, Poland) and antibiotics: penicillin (100 U/ml), streptomycin (100 μg/ml). Cells were incubated for seven days at 32°C, washed thrice with 1 × PBS, and collected for RNA isolation in Fenozol reagent (A&A Biotechnology, Poland). All research involving the infectious material was carried out adhering to the biosafety regulations. All research involving genetic modifications was carried out adhering to the national and international regulations.

### Lentivirus production and transduction

293T cells were seeded on 10 cm^2^ dishes, cultured for 24 h at 37°C with 5% CO_2_ and transfected with psPAX, pMD2G and third transfer plasmid (pWPI/KLK13, pLKO.1-TRC/shrnaKLK13 or Lego-G2) using polyethyleneimine (Sigma-Aldrich, Poland). psPAX (Addgene plasmid # 12260) and pMD2G (Addgene plasmid # 12259) was a gift from Didier Trono. pLKO.1 - TRC cloning vector was a gift from David Root (Addgene plasmid # 10878) (*94*). Cells were further cultured for 96 h at 37°C with 5% CO_2_ and lentiviral particles were collected every 24 h and stored at 4°C. Lentivirus stocks were concentrated 25-fold using centrifugal protein concentrators (Amicon Ultra, 10-kDa cutoff; Merck, Poland) and stored at - 80°C.

RD cells were seeded in T75 flasks, cultured for 24 h at 37°C with 5% CO_2_ and transduced with lentiviral particles harboring KLK13, TMPRSS2 gene or control vector in the presence of polybrene (4 µg/ml; Sigma-Aldrich, Poland). Cells were further cultured for 72 h at 37°C with 5% CO_2_ and positively transduced cells were selected using blasticidin (2 µg/ml Sigma-Aldrich, Poland). Primary human epithelial cells seeded on 10 cm^2^ dishes were cultured in BEGM medium and transduced with lentiviral particles harboring shRNA against KLK13 (a set of 3) or GFP gene in the presence of polybrene (5 µg/ml; Sigma-Aldrich, Poland). Cells were further cultured for 72 h at 37°C with 5% CO_2_ and positively transduced cells were selected using puromycin (5 µg/ml Sigma-Aldrich, Poland). Selected cells were plated on insert supports and further cultured in ALI in the presence of puromycin (1 µg/ml).

### Pseudoviruses

293T cells were seeded on 6-wells plates, cultured for 24 h at 37°C with 5% CO_2_ and transfected using polyethyleneimine (Sigma-Aldrich, Poland) with the lentiviral packaging plasmid (psPAX), the VSV-G envelope plasmid (pMD2G) or HCoV-HKU1 S glycoprotein (pCAGGS-HKU1-S) and third plasmid encoding luciferase (pRR Luciferase). pRR Luciferase was a gift from Paul Khavari (Addgene plasmid # 120798) (*97*). Cells were further cultured for 72 h at 37°C with 5% CO_2_ and pseudoviruses were collected every 24 h and stored at 4°C.

RD cells were seeded in 48-wells plates, cultured for 24 h at 37°C with 5% CO_2_ and transduced with pseudoviruses harboring VSV-G or S-HKU1 proteins or lacking the fusion protein (Δ Env) in the presence of polybrene (4 µg/ml; Sigma-Aldrich, Poland). HAE cultures were washed thrice with 100 μl of 1 × PBS and subsequently inoculated with S-HKU1 or VSV-G pseudoviruses. After 4 h incubation at 37°C unbound virions were removed by washing with 100 μl of 1 × PBS and HAE cultures were cultured at an air-liquid interphase. Cells were further cultured for 72 h at 37°C with 5% CO_2_ and lysed in luciferase substrate buffer (Bright-Glo; Promega, Poland). Lysates were transferred onto white 96-wells plates and luciferase levels were measured on a microplate reader Gemini EM (Molecular Devices, UK).

### Isolation of nucleic acids and reverse transcription (RT)

Viral DNA/RNA Kit (A&A Biotechnology, Poland) was used for nucleic acid isolation from cell culture supernatants, according to the manufacturer’s instructions. Cellular RNA was isolated using Fenozol reagent (A&A Biotechnology, Poland), followed by DNase I treatment (Thermo Fisher Scientific, Poland). cDNA samples were prepared with a High Capacity cDNA Reverse Transcription Kit (Thermo Fisher Scientific, Poland), according to the manufacturer’s instructions.

### PCR

Human KLKs mRNA was reverse transcribed and amplified in a reaction mixture containing 1 × Dream *Taq* Green PCR master mix (Thermo Fisher Scientific, Poland) and appropriate primers (**Table 3**; 500 nM each). β-actin was used as a household gene reference. The reaction was carried out according to the scheme: 5 min at 95°C, followed by 35 cycles of 30 s at 95°C, 20 s at 59°C and 20 s at 72°C, followed by 10 min at 72°C.

**Table 3.**
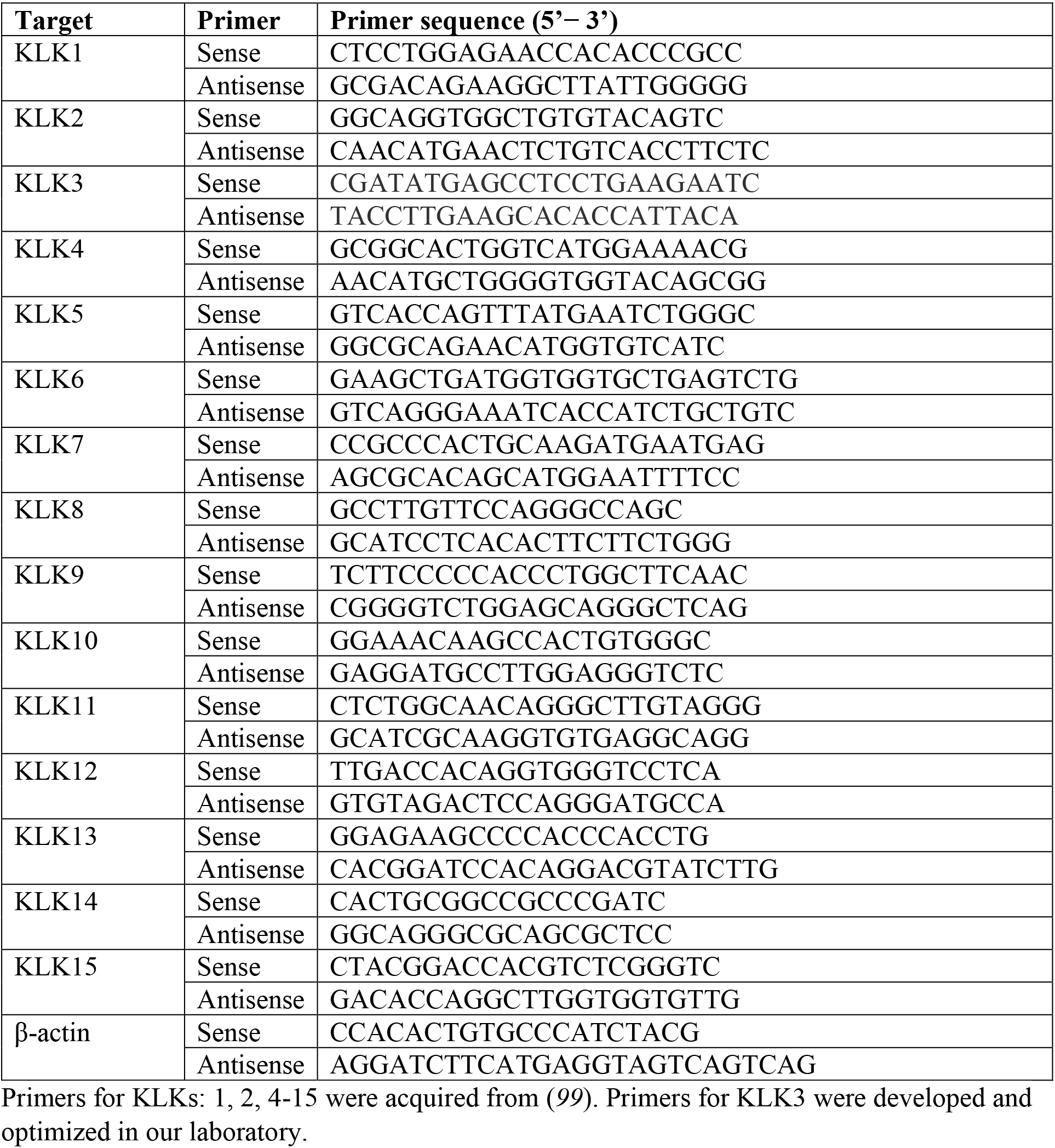
Primers used for PCR of each KLK gene.

### Quantitative PCR (qPCR)

HCoV-HKU1 RNA yield was assessed using real-time PCR (7500 Fast Real-Time PCR; Life Technologies, Poland). cDNA was amplified in a reaction mixture containing 1 × TaqMan Universal PCR Master Mix (Thermo Fisher Scientific, Poland), in the presence of FAM / TAMRA (6-carboxyfluorescein / 6-carboxytetramethylrhodamine) probe (100 nM; 5’ – TTGAAGGCTCAGGAAGGTCTGCTTCTAA– 3’) and primers (450 nM each; forward: 5’ – CTGGTACGATTTTGCCTCAA – 3’ and reverse: 5’-ATTATTGGGTCCACGTGATTG– 3’) (*98*). The reaction was carried out according to the scheme: 2 min at 50°C and 10 min at 92°C, followed by 40 cycles of 15 s at 92°C and 1 min at 60°C.

### Detection of HCoV-HKU1 N sg mRNA

Total nucleic acids were isolated from virus or mock-infected cells at 7 days p.i. using Fenozol reagent (A&A Biotechnology, Poland), according to the manufacturer’s instructions. Reverse transcription was performed using a high-capacity cDNA reverse transcription kit (Life Technologies, Poland), according to the manufacturer’s instructions. Viral cDNA was amplified in a 20 µl reaction mixture containing 1 × Dream *Taq* Green PCR master mix (Thermo Fisher Scientific, Poland), and primers (500 nM each). The following primers were used to amplify HCoV-HKU1 subgenomic mRNA (sg mRNA): common sense primer (leader sequence), 5 – TCTTGTCAGATCTCATTAAATCTAAACT-3’; nucleocapsid antisense for 1^st^ PCR, 5’ – AACTCCTTGACCATCTGAAAATTT – 3’; nucleocapsid antisense for nested PCR, 5’ – AGGAATAATGTGGGATAGTATTT – 3’. The conditions were as follows: 3 min at 95°C, 35 cycles (30 cycles for nested PCR) of 30 s at 95°C, 30 s at 49°C, and 20 s at 72°C, followed by 5 min at 72°C and 10 min at 4°C. The PCR products were run on 1% agarose gels (1Tris-acetate EDTA [TAE] buffer) and analyzed using molecular imaging software (Thermo Fisher Scientific, Poland).

### Detection of TMPRSS2 protease

After blasticidin selection, RD cells expressing TMPRSS2 (RD_TMPRSS2), KLK13 (RD_KLK13) or control cells (RD_ctrl) were scraped and collected by centrifugation. Cells were lysed in RIPA buffer (50 mM Tris, 150 mM NaCl, 1% Nonidet P-40, 0.5% sodium deoxycholate, 0.1% SDS, pH 7.5), boiled for 5 min, cooled on ice, and separated on 10% polyacrylamide gel alongside dual-color Page Ruler Prestained Protein size markers (Thermo Fisher Scientific, Poland). The separated proteins were then transferred onto a Westran S PVDF membrane (GE Healthcare, Poland) by wet blotting (Bio-Rad, Poland) for 1 h, 100 Volts in transfer buffer (25 mM Tris, 192 mM glycine, 20% methanol) at 4°C. The membranes were blocked by overnight incubation at 4°C in TBS-Tween (0.1%) buffer supplemented with 5% skimmed milk (BioShop, Canada). A mouse monoclonal anti-TMPRSS2 antibody (clone P5H9-A3; 1:500 dilution; Sigma-Aldrich, Poland), followed by incubation with a horseradish peroxidase-labeled anti-mouse IgG (65 ng/ml; Dako, Denmark) diluted in 5% skimmed milk / TBS-Tween (0.1%). The signal was developed using the Pierce ECL Western blotting substrate (Thermo Scientific, Poland) and visualized using the ChemiDoc Imaging System (Bio-Rad, Poland).

### Expression and purification of KLK13 and KLK14

The proKLK13 gene was amplified using cDNA obtained from HAE cultures and specific primers. The codon-optimized proKLK14 gene was custom-synthesized (Thermo Scientific, Poland). The products were cloned into the pLEXSY_I-blecherry3 plasmid (Jena Bioscience, Germany) and the resulting constructs were verified by sequencing. All preparations for transfection, selection, and expression of the host *Leishmania tarentolae* strain T7-TR were performed according to the Jena Bioscience protocol for inducible expression of recombinant proteins secreted to medium (LEXSinduce Expression kit, Jena Bioscience, Germany). Expression of proKLK13 and proKLK14 was induced with 15 µg/ml of tetracycline (BioShop, Canada) and carried out for 3 consecutive days. Culture media were collected and precipitated with 80% ammonium sulfate, spun down at 15,000 × g for 30 min at 4°C. Pellets were suspended in 10mM sodium phosphate pH 7.5 and dialyzed overnight at 4°C into 10mM sodium phosphate pH 7.5. The KLKs were isolated *via* the 6 × His tag using nickel resin (GE Healthcare, Poland) according to the manufacturer’s protocol. Obtained fractions were analyzed by SDS PAGE in reducing conditions and fractions containing proKLK13 or proKLK14 were concentrated with Vivaspin® 2 (Sartorius, Germany) and further purified using size exclusion chromatography (Superdex s75 pg; GE Healthcare, Poland). Fractions containing proKLK13 or proKLK14 were concentrated and the buffer was changed to 50mM Tris pH 7.5, 150 mM NaCl. After purification and self-activation at 37°C for 24 h, activity of proteases was assessed by serine protease inhibitor Kazal-type 6 (SPINK6) titration, as described previously (*63*).

### Cloning of HmuY-based CleavEx fusion proteins

The fusion constructs were based on positions 26-216 of the *Porphyromonas gingivalis* HmuY protein-encoding gene (accession number ABL74281.1), employed as a carrier protein. The gene was amplified using Phusion DNA polymerase (Thermo Scientific, Poland) and specific primers (forward: 5’– ATATGCGGCCGCAGACGAGCCGAACCAACCCTCCA – 3’ and reverse: 5’ – ATACTCGAGTTATTTAACGGGGTATGTATAAGCGAAAGTGA – 3’) from whole-genomic DNA isolated from *Porphyromonas gingivalis* strain W83. PCR conditions were as follows: 98°C for 30s, followed by denaturation at 98°C for 10s, annealing at 68°C for 40s and extension at 72°C for 30s/kb over 35 cycles with a final extension at 72°C for 7 min. Then, the HmuY gene was further amplified in the three consecutive PCR reactions with primers specific to the 5’ HmuY fragment and 3’-specific primer introducing additional nucleotides dependent on the designed sequence (proKLK13 primers, **Table 4**). The reaction was ligated into a modified pETDuet plasmid (potential tryptic cleavage sites were removed from the MCS) using Quick Change mutagenesis using Phusion DNA polymerase (Thermo Scientific, Poland). Alternatively, the designed fusion protein-encoding sequences were produced using Phusion Site-Directed Mutagenesis (Thermo Scientific, Poland), *via* sequence exchange of the previously prepared CleavEx construct (HKU1-S primers, **Table 4**). The final product was transformed into competent *E. coli* T10 cells and further purified and sequenced.

**Table 4.**
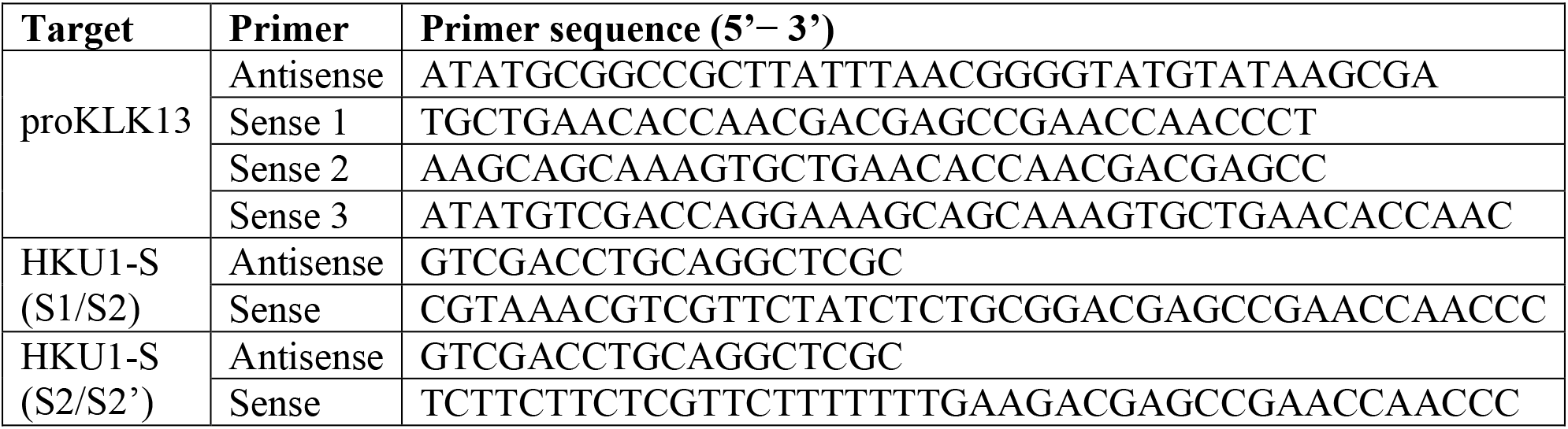
Primers used in the CleavEx design.

### Expression and purification of CleavEx fusion proteins

Protein expression was performed in *E. coli* BL21 and was induced by the addition of 0.5 mM IPTG to the bacterial culture (OD_600_ 0.5-0.6), followed by shaking for 3h at 37°C. Then, the bacteria were spun down and the pellet was suspended in buffer A (10 mM sodium phosphate, 500 mM NaCl and 5 mM imidazole, pH 7.4). The pellet suspension was then sonicated and spun down. Soluble proteins were purified using HisTrap^TM^ Excel (GE Healthcare, Poland) column in buffer A with a linear gradient of 0-100% of 1 M imidazole in buffer A in 20 column volumes. Fractions containing protein of interest were pooled together and the buffer was exchanged to 50 mM Tris pH 7.5. Lastly, the protein of interest was purified by ion-exchange chromatography using MonoQ 4.6/100 PE column (GE Healthcare, Poland) with a linear gradient of 0-100% 50 mM Tris pH 7.5, 1M NaCl in 15 column volumes.

### Expression and purification of the HCoV-HKU1 Spike protein

293T cells were seeded on 60 cm^2^ dishes, cultured for 24 h at 37°C with 5% CO_2_ and transfected with 25 μg of pSecTag2-HKU-S plasmid per dish using polyethyleneimine (Sigma Aldrich, Poland). Cells were further cultured for 72 h at 37°C with 5% CO_2_ and collected for HKU1-S purification. Cell pellets were lysed in RIPA buffer (50 mM Tris, 150 mM NaCl, 1% Nonidet P-40, 0.5% sodium deoxycholate, 0.1% SDS, pH 7.5) in the presence of Viscolase (1250 U/ml; A&A Biotechnology, Poland), clarified by centrifugation, and filtered (0.45 µm syringe PES filter). Supernatant containing 6 × His tagged S protein was mixed in 1:2 ratio with binding buffer (20 mM NaH_2_PO_4_, 500 mM NaCl, 20 mM imidazole, pH = 7.4) and purified using a fast performance liquid chromatography system (FPLC; AKTA, GE Healthcare, Poland) with a Ni^2+^ HiTrap IMAC (2 × 1 ml) column (GE Healthcare, Poland) preequilibrated with the binding buffer. The 6 × His tagged S protein was eluted with elution buffer (20 mM NaH_2_PO_4_, 500 mM NaCl, 500 mM imidazole, pH = 6.9). The control sample from mock-transfected cells was prepared in the same manner. Fractions containing 6 × His tagged S protein or the respective fractions from control purification were pooled and dialyzed against phosphate-buffered saline (PBS) with 5% glycerol.

### CleavEx screening assay and HKU1-S cleavage

A total of 15 ng of each CleavEx protein was incubated with 50, 250 and 500 nM KLK13, respectively in 50 mM Tris pH 7.5. For the full-length Spike protein, fractions containing purified HKU1-S or mock samples were incubated with 0.5, 1.0 or 5.0 μM KLK13, respectively in 50 mM Tris pH 7.5. Samples were incubated at 37°C for 3 hours and immediately halted with the addition of 50 mM DTT-supplemented SDS sample buffer (1:1), boiled for 5 min, cooled on ice, and separated on 10% polyacrylamide gels alongside dual-color Page Ruler Prestained Protein size markers (Thermo Fisher Scientific, Poland). The separated proteins were then transferred onto a Westran S PVDF membrane (GE Healthcare, Poland) by wet blotting (Bio-Rad, Poland) for 1 h, 100 Volts in transfer buffer: 25 mM Tris, 192 mM glycine, 20% methanol at 4°C. The membranes were then blocked by overnight incubation (at 4°C) in TBS-Tween (0.1%) buffer supplemented with 5% skimmed milk (BioShop, Canada). A horseradish peroxidase-labeled anti-His tag antibody (1:25000 dilution; Sigma-Aldrich, Poland) diluted in 5% skimmed milk / TBS-Tween (0.1%) was used to detect the His-tagged HmuY proteins. The signal was developed using the Pierce ECL Western blotting substrate (Thermo Scientific, Poland) and visualized using the ChemiDoc Imaging System (Bio-Rad, Poland).

### Identification of the cleavage site

A total of 10 µg S1/S2 CleavEx protein was incubated with 500 nM of KLK13 at 37°C for 5 h. Next, the reaction was stopped by the addition of 50 mM DTT-supplemented SDS sample buffer (1:1) and samples were immediately boiled for 5 min. The samples were then separated on 10% polyacrylamide gel alongside the dual-color Page Ruler Prestained Protein size markers (Thermo Fisher Scientific, Poland) in the Tris-Tricine SDS-PAGE system. The separated proteins were then electrotransferred onto a Western S PVDF membrane (GE Healthcare, Poland) using the Trans-Blot SD Semi-Dry Transfer Cell (Bio-Rad, Poland). The transfer was performed for 30 min at 15 V in transfer buffer (10 mM N-cyclohexyl-3-aminopropanesulfonic acid, 10% methanol, pH 11). Following the transfer, the membrane was stained with 0.025% (w/v) Coomassie Brilliant Blue R-250 (BioShop, Poland) and the bands of interest were sequenced *via* Edman degradation using a PPSQ-31A automatic protein sequencer (Shimadzu, Japan).

## ACKNOWLEDGEMENTS

This work was supported by grants from the National Science Center UMO-2013/08/W/NZ1/00696 (J.P.), UMO-2016/22/E/NZ5/00332 (T.K.), UMO-2013/08/S/NZ6/00730 (A.N.), UMO-2012/07/E/NZ6/01712 and UMO-2017/27/B/NZ6/02488 (K.P.). Faculty of Biochemistry, Biophysics and Biotechnology of the Jagiellonian University is a partner of the Leading National Research Center supported by the Ministry of Science and Higher Education of the Republic of Poland. The authors are grateful to Xingchuan Huang for providing pCAGGS/HKU1-S plasmid, Ambra Saraccino for providing pWPI vector, Laura Sasiadek for providing KLK primers used in the study and Grzegorz Dubin for providing reference samples. The funders had no role in study design, data collection, and analysis, decision to publish, or preparation of the manuscript.

## AUTHOR CONTRIBUTIONS STATEMENT

A.M., K.F., E.B., M.K, A.N. and P.M conducted the experiments. A.L., T.K., M.O., M.U. and J.P. provided materials and methods for the study. A.M. and K.P. designed the study and experiments, analyzed the data, and wrote the manuscript. K.P. and T.K. supervised the study. All authors reviewed the manuscript and approved the submitted version. All authors agreed to be personally accountable for their contributions and to ensure that questions related to the accuracy or integrity of any part of the work are appropriately investigated, resolved, and the resolution documented in the literature.

